# Urine microbiome changes during and after radiotherapy for prostate cancer

**DOI:** 10.1101/2024.04.15.589478

**Authors:** Michał Złoch, Ewelina Sibińska, Fernanda Monedeiro, Wioletta Miśta, Adrian Arendowski, Piotr Fijałkowski, Monika Pietrowska, Jolanta Mrochem-Kwarciak, Anna Jędrzejewska, Ewa Telka, Kinga Karoń, Małgorzata Rabsztyn, Paweł Pomastowski, Dorota Gabryś

## Abstract

**Background:** The urinary microbiome may play a new important role in the development of complications, but still, there is no information about their changes during and after radiotherapy (RT). This study aimed to use the matrix-assisted laser desorption/ionization mass spectrometry (MALDI MS) technique to identify the microbiome and assess its changes in urine samples of 88 patients irradiated for prostate cancer.

**Material and methods:** Blood for biochemical analysis and urine samples for MALDI were collected at various time points before gold fiducial implantation (t1) at the beginning (t2) and end of radiotherapy (t3); during follow-up, 1 (t4), 4 (t5), 7 (t6) months after the end of treatment.

**Results:** We identified 1801 different microbial isolates, in 89% (470/528) samples revealed the presence of at least one microbial species among which 79% (373/470) were polymicrobial. Species level: 136 G+, 29 G-, 2 *Candida* have been noted. The far most abundant group of the identified microorganisms was *Staphylococcus* members −51.6% of all isolates followed by *Micrococcus* (9.1%), *Enterococcus* (7.6%), *Kocuria* (5.6%), *Corynebacterium* (5.4%), and *Streptococcus* (2.2%). A lower variety of microorganisms incident was observed at the end of RT. The total number of species (TNS) was 50 at t1, increased up to 61 at t2, and then fell to the initial value of 52 at t3. The increase in biodiversity was noted after radiotherapy t4-68, t5-86, and t6-75 (p<0.05). Changes in the biodiversity of the urinary microbiota were also reflected in the differences in the total number of isolates (TNI) – 261, 281, and 273 for time points t1-t3 compared to the 292, 362, and 332 for time points t4-t6 as well as in the total number of detected genera (TNG) – 25, 29, 23 (t1-t3) and 28, 38, 31 (t4-t6). *Actinomyces*, *Corynebacterium*, *Staphylococcus*, *Streptococcus*, demonstrated significant correlation with the RT stages. Concerning individual species, only *K. rhizophila* abundance significantly increased with time (p=0.045). Bacteria incidence was strongly correlated with glucose levels in urine. The same correlation was observed for glucose levels in blood, but in a weak manner. Staphylococcus presence was related to higher tPSA.

**Conclusion:** RT for prostate cancer induces a dynamic response in the urinary microbiome, characterized by an initial reduction in diversity post-RT followed by a subsequent increase. Our findings highlight the significant influence of glucose levels in both urine and blood on the urinary microbiota. These insights contribute to the evolving understanding of the interplay between RT, the urinary microbiome, and patient health, paving the way for more targeted interventions and personalized approaches in prostate cancer treatment.

## 1. INTRODUCTION

Modern radiotherapy technologies allow for precise irradiation in the lower abdomen and pelvis such as prostate cancer, bladder cancer, cervical cancer, anal cancer, limiting the impact on surrounding tissues(^1^). But still, radiotherapy not only treats the cancer but also irradiates nearby other anatomical structures such as the urinary tract or intestines, where some early and late side effects are often observed(^2,3^). Some of our patients report urinary tract problems which depend on the dose, on the amount of healthy tissue that is exposed to the radiation, and they are patient specific(^4^). Urinary toxicity may decrease patients quality of life presenting as difficult, frequent, or painful urination, urinary leakage, blood in urine, abdominal cramping, and nocturia. Radiotherapy induced toxicity is a well-known fact after the treatment of prostate cancer and potential mechanisms are also determined. Nevertheless, the urinary microbiome may play a new important role in the development of complications.

It is now well-known that urine is not sterile, and it holds a wide variety of microbial species, identified as the urinary microbiota(^5^). The acknowledgment of a thriving urinary microbiota challenges the long-standing assumption of sterility in the urinary tract of healthy individuals, emphasizing the dynamic interplay between microorganisms and urological health. Recognizing the urinary environment as a complex ecosystem opens avenues for innovative diagnostic and therapeutic strategies, offering a glimpse into a future where urologists consider and leverage the microbiome for a more comprehensive understanding and management of urinary conditions(^6^). Besides its role in tumorigenesis and cancer progression beyond its role, can be used as a new potential biomarker in the diagnosis, prognosis, and risk stratification of this disease(^7^).

Alterations in the microbiome have the potential to serve as indicators for predicting disease progression and treatment outcomes. Nearly all microbial-based cancer diagnosis is sequencing-based and focus on gastrointestinal cancers. Many of these studies have shown that the microbiome contributes significantly to the development of certain types of cancer; in particular, the contribution of the fecal microbiome to gastrointestinal cancers(^8^). Moreover, the specific gut bacterial community profiles are associated with response to systemic therapies such as chemotherapy(^9^). Only recently has it been suggested that other types of cancer (such as head and neck cancers(^10^), lung cancer(^11,12^), skin cancer(^13^), pancreatic cancer(^14^)) may also harbor microbiota with unique compositions. Prostatic cancer is not an exception and many studies reported associations between certain patterns of urinary microbiome and prostate cancer(^15–17^). Bacteria have been widely considered as potential contributors to chronic, low-grade inflammation, a factor that could potentially lead to tumor development. It is challenging to assert that a specific microorganism has a substantial impact on the development of cancer due to the intricate and frequently unknown interactions in microbial epidemiology.

To date, most of the studies devoted to examining the impact of the human microbiota on the outcomes of prostate cancer treatment, i.e., via radiation therapy, were focused on investigating prostate tumor microenvironment (biopsy) or gut microflora composition(^18^). Nevertheless, advances in molecular biology techniques and culture methods have made it possible to define the urinary microbiome associated with urine most commonly collected from midstream(^19–21^). Although there is evidence that urinary microbiota is implicated in prostatic diseases, better delineation of its role in both prostate cancer development as well as its treatment progression is still needed(^5^).

This study aimed to use the MALDI technique with previously established conditions(^4^) to identify the microbiome and assess its changes in urine samples of patients irradiated for prostate cancer. The composition of the microbiome was analyzed at various time points before gold fiducial implantation to assess the primary bacterial composition in urine (before the use of antibiotics); then at the beginning and end of radiotherapy; and during follow-up, 1, 4, 7 months after the end of irradiation. Moreover, changes in the species composition of microorganisms were correlated with changes in blood tests and basic urine tests.

## 2. MATERIALS AND METHODS

### 2.1. Patients’ characteristic

Patients who were irradiated primarily for prostate cancer at all stages, with or without hormonal treatment, were included in the study. Before undergoing planning procedures for radiotherapy, all patients had one to three gold fiducials implanted. Radiotherapy planning, radiation treatment and post-treatment follow up were carried out at one center the National Institute of Oncology. Maria Skłodowska Curie in accordance with the applicable protocol. Patients characteristic are shown in **Tab. 1**.

**Tab. 1.** Patient characteristics.

| Patient Characteristics | N / % |
| --- | --- |
| Age median 69 (range 50-88) |  |
| Clinical stage |  |
| T1a-T1c | 36 / 41.91% |
| T2a-T2c | 40 / 45.45% |
| T3-T4 | 12 / 13.64% |
| PSA |  |
| <10 | 42 / 47.73% |
| 10–20 | 25 / 28.41% |
| >20 | 19 / 21.59% |
| No data | 2 / 2.27% |
| Gleason score |  |
| 6 | 34 / 38.64% |
| 7 (3 + 4) | 16 / 18.18% |
| 7 (4 + 3) | 16 / 18.18% |
| 8 | 16 / 18.18% |
| 9–10 | 6 / 6.82% |
| Use of androgen deprivation therapy |  |
| Yes | 67 / 76.14% |
| No | 21 / 23.86% |
| Treatment |  |
| Fractionated Linear accelerator | 45 / 51.14% |
| SBRT Linear accelerator | 4 / 4.45% |
| SBRT Cyberknife | 39 / 44.32% |
| Irradiated volume |  |
| Prostate alone | 51 / 57.95% |
| Prostate bed + pelvic lymph nodes | 1 / 1.14% |
| Prostate + pelvic lymph nodes | 36 / 40.91% |

As previously reported the treatment decision was made according to disease stage, histopathological grade, and PSA level. Participation in the study did not affect the choice of method, irradiated volumes, and dose. Majority of patients (76%) were on hormonal treatment as analog LH-RH, with or without Flutamide. Radiotherapy was delivered on a linear accelerator, or cyberknife. The volumes included: prostate alone, prostate with a base of seminal vesicles, prostate with the seminal vesicles, irradiated to a total dose of 76–78 Gy delivered in 2 Gy/fraction standard fractionation, or hypofractionation of 36.25 Gy delivered in 7.25 Gy/fraction, using MV photons. Two patients had a prostate boost of 15 Gy delivered with brachytherapy. If required, pelvic lymph nodes were irradiated to a total dose of 44–50 Gy delivered in 2 Gy/fraction using MV photons, with or without a boost to the involved lymph nodes to a total dose of 60–68 Gy, or delivered as a stereotactic boost of 16 Gy in 2 fractions. Appropriate regulatory approval, including ethical approval, was obtained in all jurisdictions. The study was approved by the NIO-PIB Ethics Committee KB/430-104/19 and conducted in accordance with the principles of Good Clinical Practice. All patients provided written informed consent.

### 2.2. Urine and blood samples

Urine and blood samples were systematically collected from 88 prostate cancer patients who undergo radiotherapy, a total of 528 urine samples were obtained. The sample collection process was executed at six stages throughout their treatment course, denoted as follows: Stage t1 – before the placement of gold fiducial markers within the prostate gland, Stage t2 – on the first day of radiotherapy, Stage t3 – on the last day of radiotherapy, Stage t4 – one month after radiotherapy, Stage t5 – four months after radiotherapy, and Stage t6 – seven months after radiotherapy as previously reported(4). At time point t1, patients had blood and urine collected at least 1 day before starting antibiotics and before rectal debridement. Usually, Azithromycin 500 mg or Ciprofloxacin 250 mg was prescribed. An enema infusion the evening before and the morning of the day of gold marker implantation was usually recommended for cleaning the rectum before the procedure.

Urine samples were collected from midstream into sterile containers in duplicate. One container was delivered to the hospitals diagnostic laboratory for routine tests, while the second one was frozen at −80°C and subsequently sent to the Centre for Modern Interdisciplinary Technologies in Toruń for microbiome identification using the MALDI technique.

### 2.3. Isolation and culturing of microorganisms

Following the defrosting process at room temperature and thorough vortexing, the urine samples were directly inoculated onto five solid culture media: Columbia Blood Agar (BLA; Oxoid, Basing-stoke, Great Britain) with 5% (v/v) sheep blood (GRASO Biotech, Starogard Gdański, Poland), CHROMagar Orientation (CHRA; GRASO Biotech, Starogard Gdański, Poland), Glucose Bromocresol Purple Agar (BCP; Sigma Aldrich, Steinheim, Germany), CLED Aagr (CLED; Sigma Aldrich, Steinheim, Germany), and Schaedler Agar (SCH; Sigma Aldrich, Steinheim, Germany). For this purpose, 100 µl of urine was applied to the substrate and evenly spread using a spatula. Cultures were conducted under aerobic conditions for samples on BLA, BCP, CLED, and CHRA media, while cultures in an environment with an increased CO_2_ content of up to 5% were carried out for SCH medium. All samples were incubated at 37 °C for a duration of 40 to 48 hours. To achieve pure cultures, individual colonies with distinct morphologies were aseptically transferred to fresh plates, streaked, and then incubated at 37 °C for 18 to 24 hours.

### 2.4. MALDI identification

All microorganism isolates were identified using the direct colony extraction method on a plate. A freshly grown colony from an overnight culture was smeared onto a 96-spot steel target plate and coated with 1 μL of 70% formic acid (Sigma-Aldrich). Each spot was allowed to dry before being overlaid with 1 µL of α-cyano-4-hydroxycinnamic acid (CHCA) matrix solution (Bruker Daltonics). After complete drying, the target plate was inserted into the MALDI mass spectrometer.

Measurements were automatically conducted utilizing the Microflex LT MALDI-TOF MS system (Bruker Daltonik GmbH), employing a positive linear mode. The FlexControl software integrated with MBT Compass version 4.1 was used in conjunction with a 60 Hz nitrogen laser operating at a wavelength of 337 nm. The instrument was routinely calibrated, following the manufacturer’s recommendation, using the bacterial test standard (BTS, Bruker Daltonics). Mass spectra were recorded within the mass-to-charge ratio (m/z) range of 2,000 to 20,000. Each spectrum underwent interpretation facilitated by the MALDI Biotyper automation control, utilizing the Bruker Biotyper 4.1 software and library (version H 2021, which comprised 10,834 entries). The accuracy of identification was assessed using a scoring system provided by the manufacturer. Identification scores below 1.700 signified no successful identification, scores falling within the range of 1.700 to 1.999 indicated identification at the genus level, and scores exceeding 2.000 indicated identification down to the species level.

If accurate identification was not achieved for certain isolates, the test was repeated using the method involving the preparation of protein extracts with acetonitrile and formic acid. The specific procedure for this method is outlined in our prior publication(4).

### 2.5. Statistical analysis

Data analysis was performed in R environment (R v.4.2.1), using RStudio console (v. 2022.02.03, PBC, Boston, MA, USA). A dot plot representing the total incidence of microbial genera per time point was prepared using “ggplot2” package. Networks showing concurrent microorganisms detected in samples, at each time point, were built using “sna” package. Hierarchical cluster analysis was conducted to identify microbiome composition profiles per assessed time point, for this purpose gplots::heatmap.2 was used, applying Spearman rank correlation as agglomeration method. To assess the dependence between the incidence of microbial species detected in urine samples and the different time points referring to the patient’s treatment course, a chi-square test was performed using stats::chisq.test function. In case the assessed expected frequencies were less than 5, Fisher’s exact test was performed instead, using “stats::fisher.test”. Correlations between the incidence of detected microbial species, assessed time points and further biochemical parameters were evaluated using “Hmisc” package, employing Spearman’s method. A correlation plot depicting these results were prepared using “corrplot” package. Significant differences between the values of biochemical parameters recorded at different time points were assessed using ANOVA (stats::aov), followed by Tukey’s Honest Significant Difference (HSD) as a post hoc test (stats::TukeyHSD). Differences between the mean values of biochemical parameters depending on the incidence of most predominant bacterial genera were evaluated using *t*-test (rstatix::t_test).

## 3. RESULTS

### 3.1. Microbial composition of the urine samples

Applied culture conditions allowed for isolation and identification 1801 different microbial isolates -9 yeasts (*Candida*) and 1792 bacteria. In 89% (470/528) of samples revealed the presence of at least one microbial species among which 79% (373/470) demonstrated a polymicrobial nature. Performed MALDI identification revealed the presence of the 55 different microbial genera in the following order – gram-positive bacteria – 36 genera (94.9% isolates), gram-negative bacteria – 18 genera (4.6% isolates), and yeasts – 1 genus (0.5% isolates) (**Fig. 1**).

**Fig. 1.**
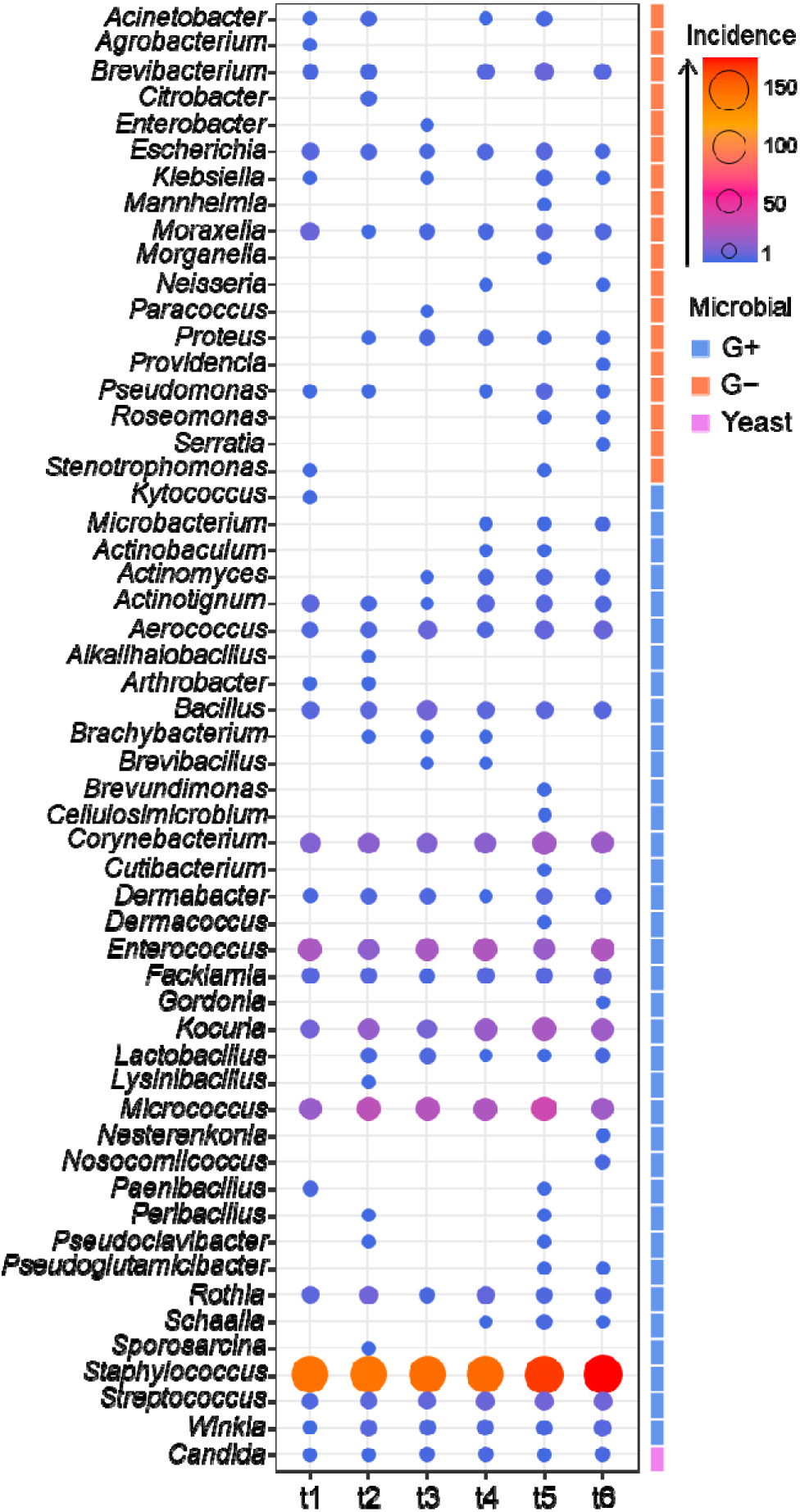
Abundance of the identified bacterial and yeast genera detected within urine samples collected from the 88 patients undergoing radiation therapy for prostate cancer. The size and color of the dots refer to detected microbial counts per genus in the respective samples. G- - gram-negative bacteria; G+ - gram-positive bacteria; value – number of isolates; t1 - before gold fiducial implantation into the prostate gland; t2 - before radiotherapy; t3 - at the end of radiotherapy; t4 - 1 month after radiotherapy; t5 - 4 months after radiotherapy; t6 - 7 months after radiotherapy.

Regarding species level, 136 gram-positive, 29 gram-negative as well as 2 *Candida* species have been noted. The far most abundant group of the identified microorganisms comprised *Staphylococcus* members - 51.6% of all isolates – represented by 24 species among which *S. epidermidis* (14%)*, S. hominis* (10.3%), and *S. haemolyticus* (8.8%) were dominant. Another relatively high abundant genus was *Micrococcus* (9.1%) followed by *Enterococcus* (7.6%), *Kocuria* (5.6%), *Corynebacterium* (5.4%), and *Streptococcus* (2.2%). The percentage of each of the other 49 genera was less than or equal to 1.6%, mostly ≤1.0% (43). The most frequently isolated gram-positive species, besides dominant *Staphylococcus* members, were *Micrococcus luteus* (8.8%), *Enterococcus faecalis* (7.4%), *Kocuria rhizophila* (4.7%), *Corynebacterium tuberculostearicum* (2.4%), *Facklamia hominis* (1.3%), *Corynebacterium glucuronolyticum* (1.2%), *Aerococcus urinae* (1.1%), *Actinotignum schaalia* (1.0%), and *Winkia neuii* (1.0%). In the case of gram-negative bacteria, *Escherichia coli* and *Moraxella osloensis* were the dominant ones – each 1.0% of all identified isolates.

### 3.2. Changes in the urinary microbiota composition during radiotherapy

Statistical analysis of the changes in the microbial composition of the urine samples were found depending on the time of sample collection during and after treatment regarding the time of the sample collection revealed the RT stage’s relevant impact on the patients’ urinary microbiome (**Fig. 2**). A lower variety of microorganisms incident was observed at the end of RT (**Fig. 2C**), where a more compact network was displayed, meaning that samples mostly comprised of core (the most dominant) bacterial genera, like *Staphylococccus*, *Enterococcus*, *Micrococcus* or *Corynebacterium*. Opposite results were depicted for time points D, E and especially for F (1, 4, and 7 months after RT) where networks were more diffused meaning more diverse species composition among samples and a higher share of less frequently isolated genera, eg. *Actinomyces*, *Paenibacillus*, *Roseomonas*, *Brevibacterium*, *Peribacillus* and so one. According to Fisher’s exact test and Pearson’s Chi-squared test, there is a significant relationship between the number of species and sampling time points (p<0.05). Regarding this, two phases of change were noted. Firstly, the total number of species (TNS), which equals 50 at the beginning of the experiment (t1), increased up to 61 after gold fiducial implantation (t2) and then fell to initial values (TNS=52) right after the end of RT (t3). Secondly, the considerable increase in biodiversity of the urinary microbiota composition was noted within 1-7 months after RT, where the peak of the increase in species incidence was observed four months after radiotherapy (t5) – 86 different species. Although TNS in samples collected one and seven months after the end of RT were lower than those collected at t5, the values were still much higher compared to the beginning of the experiment – 68 (t4) and 75 (t6). Changes in the biodiversity of the urinary microbiota were also reflected in the differences in the total number of isolates (TNI) – 261, 281, and 273 for time points t1-t3 compared to the 292, 362, and 332 for time points t4-t6 as well as in the total number of detected genera (TNG) – 25, 29, 23 (t1-t3) and 28, 38, 31 (t4-t6).

**Fig. 2.**
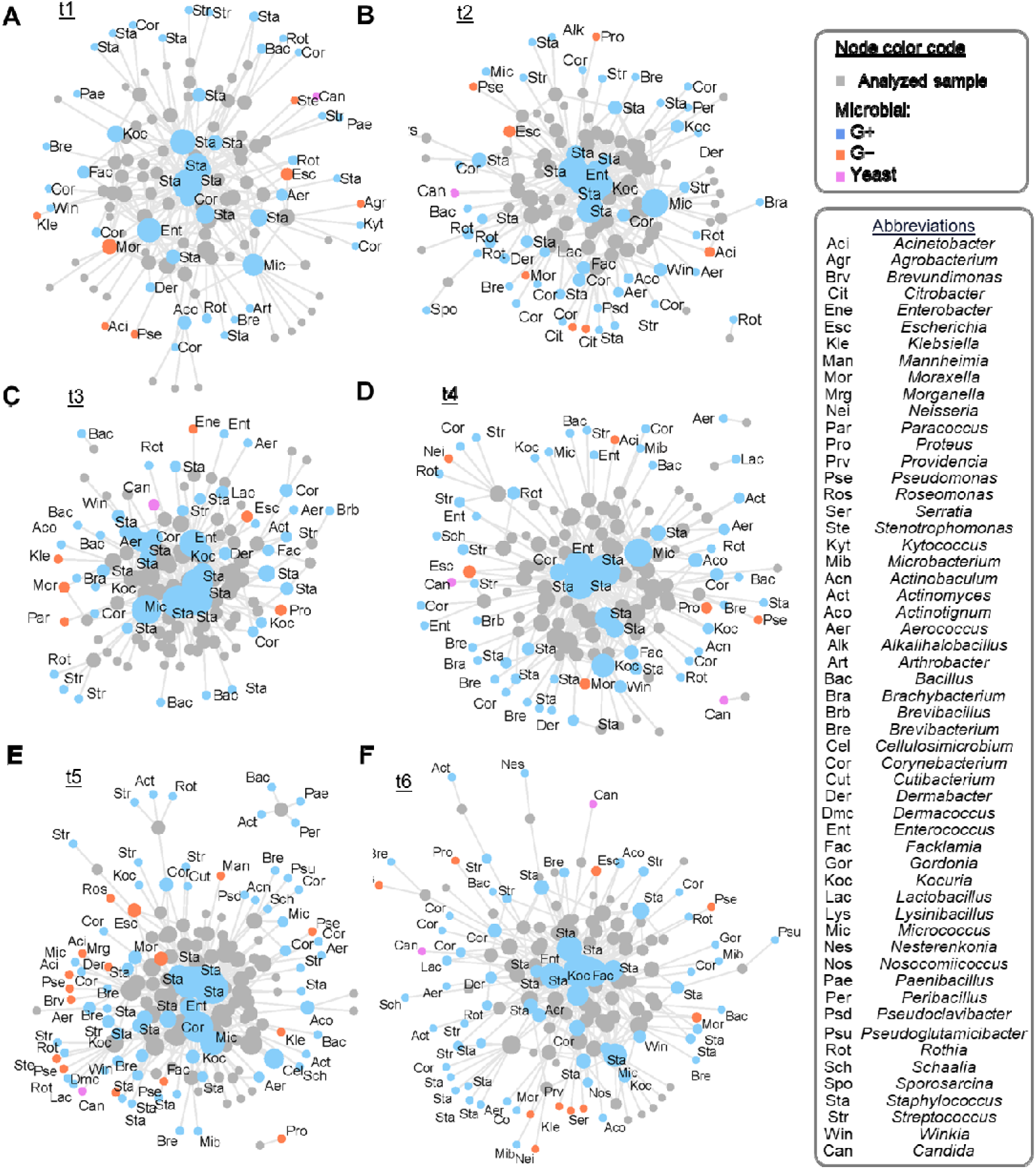
Networks depict the urinary microbiome composition at different time points. Grey nodes represent samples and other node colors refer to the microbial classification (gram-positive - blue, gram-negative - orange or yeast - pink). The size of a node is proportional to the number of links connected with this particular node. Thus, node size for microbials is proportional to the number of samples in which a given microbial was detected, and for samples it is proportional to the number of microbials detected in it. G- - gram-negative bacteria; G+ - gram-positive bacteria; A - before gold fiducial implantation into the prostate gland (t1); B - before radiotherapy (t2); C - at the end of radiotherapy (t3); D - 1 month after radiotherapy (t4); E - 4 months after radiotherapy (t5); F - 7 months after radiotherapy (t6).

Performed analysis of the correlation between the genera incidence and RT time showed that some genera displayed quantitative correlation with the passed time (**Tab. 2**). Among investigated 55 different microbial genera, seven gram-positive (*Microbcaterium*, *Actinomyces*, *Arthrobacter*, *Corynebacterium*, *Pseudoglutamicibacter*, *Staphylococcus*, and *Streptococcus*) and one gram-negative (*Roseomonas*) they have demonstrated relevant correlation with the RT stages. Only the abundance of the *Arthrobacter* genus members decreased along the passing RT time. At the same time, the rest’s quantity increased when RT passed. However, it should be mentioned that half of the mentioned genera - *Arthrobacter*, *Microbacterium*, *Pseudoglutamicibacter*, and *Roseomonas*, were inferior represented among all identified isolates - ≤ 4 isolates in total.

**Tab. 2.** The values of correlation coefficients showing genera displayed quantitative correlation with the passed time. Only significant correlations (p<0.05) are shown.

|  | *Microbacterium* | *Actinomyces* | *Arthrobacter* | *Corynebacterium* | *Pseudoglutamicibacter* | *Roseomonas* | *Staphylococcus* | *Streptococcus* |
| --- | --- | --- | --- | --- | --- | --- | --- | --- |
| **Time** | **0.92** | **0.88** | **-0.83** | **0.82** | **0.83** | **0.83** | **0.9** | **0.98** |

Concerning individual species, *Bacillus cereuş K. rhizophila*, *M. luteus*, *S. epidermidis*, and *Staphylococcus saprophyticus* can be considered as those whose incidence was more related with the RT time (**Tab. 3**). In the case of *K. rhizophila*, which abundance after 1 – 7 months after end of RT increased, this dependence was statistically significant (p=0.045).

**Tab. 3.** List of the microbial species whose incidence was correlated with the time points (Fisher’s exact test for count data with a simulated p-value based on 2000 replicates).

| **Microbial species** | **p-value** |
| --- | --- |
| *Bacillus cereus* | 0.159 |
| *Kocuria rhizophila* | 0.045* |
| *Micrococcus luteus* | 0.084 |
| *Staphylococcus epidermidis* | 0.139 |
| *Staphylococcus saprophyticus* | 0.130 |

\* - statistically relevant

Investigation of the microbial genus profiles for the different RT stages revealed the presence of the 5 clusters (**Fig. 3**).

**Fig. 3.**
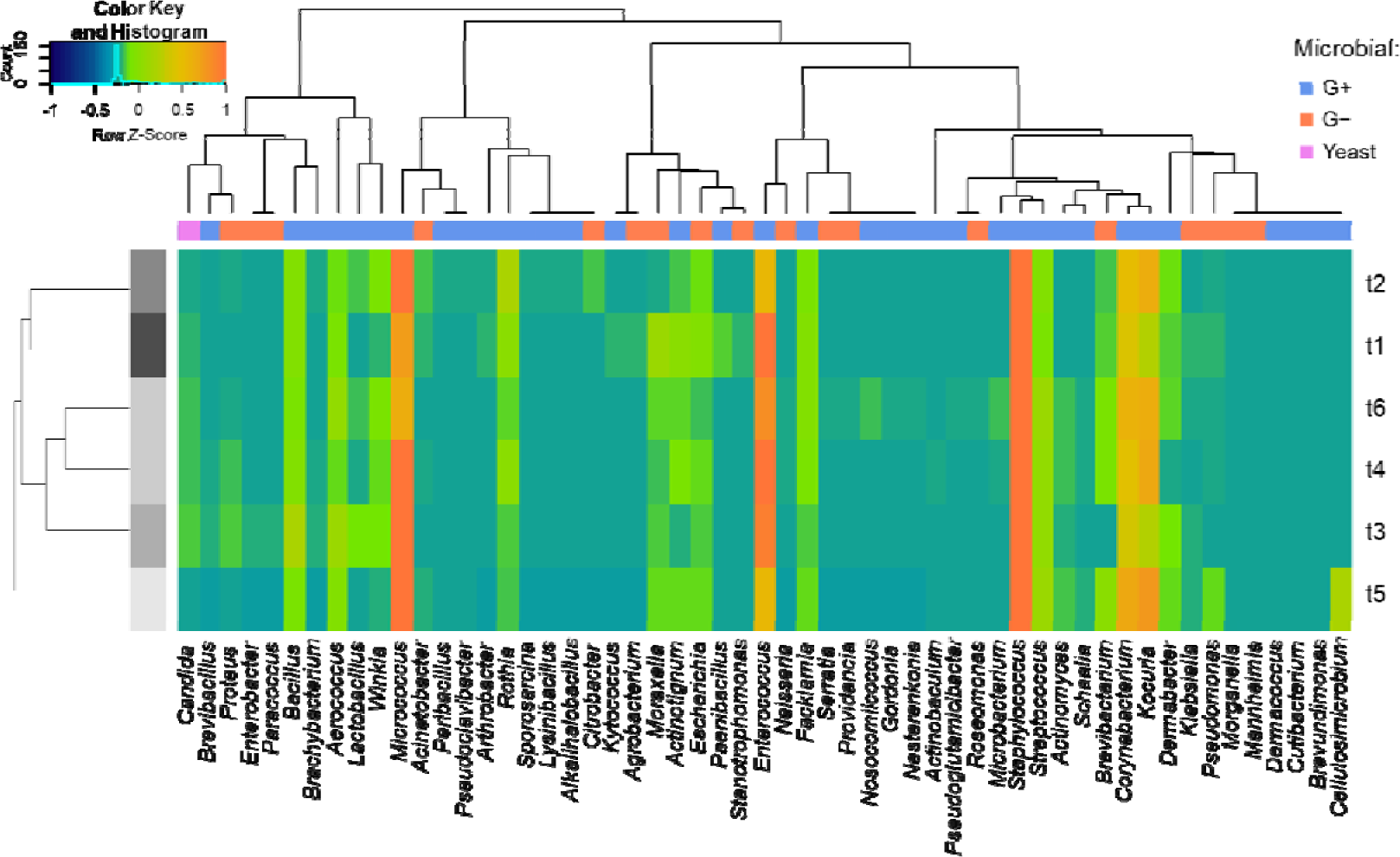
Heatmap with cluster analysis performed using Spearman rank correlation method of the time point-associated microbiota. G- - gram-negative bacteria; G+ - gram-positive bacteria; t1 - before gold fiducial implantation into the prostate gland; t2 - before radiotherapy; t3 - at the end of radiotherapy; t4 - 1 month after radiotherapy; t5 - 4 months after radiotherapy; t6 - 7 months after radiotherapy.

The analysis showed that, based on a comparison of microbiological profiles, we can divide the samples into two main groups: those collected before starting radiotherapy (t1 and t2) and those collected during and after therapy - t3 - t6. Moreover, in the latter group we can distinguish two phases of the microbial species composition change. Phase 1 – a considerable shift towards the unique microbial profile between t3 and t5 (the most distinct species pattern four months after the end of RT), which seemed to start returning to the initial composition (phase II)-one (t4) and seven months after RT (t6) were the most similar grouping in one cluster. Concerning the type of microorganisms, yeast species presented the most differentiated profiles across time points among all detected microorganisms.

### 3.3. Correlation between urinary microbiota and blood/urine biochemical parameters during radiation therapy

Performed analysis revealed the presence of dozens of significant correlations between examined parameters, including both weak, moderate, or strong positive and negative interactions (**Fig. 4**).

**Fig. 4.**
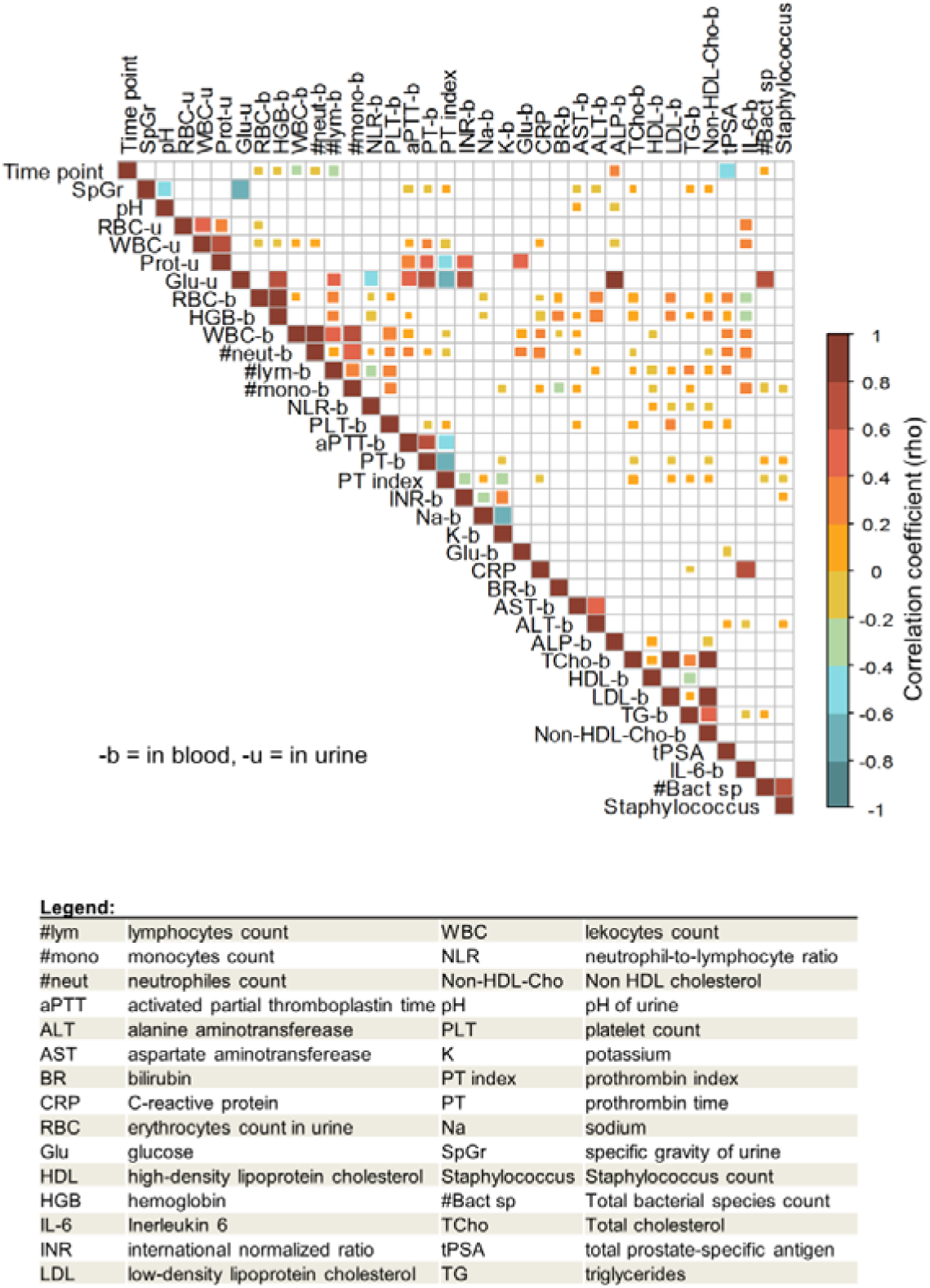
Correlation matrix demonstrating the interactions between measured urine and blood biochemical parameters including urinary microbial count for all time points (RT stages). Empty boxes – no significant correlation.

Bacteria incidence was strongly correlated (in a positive manner) with glucose level in urine. The same correlation was observed for glucose levels in blood, but in a weak manner. Weak but significant positive correlations were additionally found for bacterial species count and RT stage (time points). The incidence of *Staphylococcus* was included in the correlation analysis, because these were the most prevalent species found in urine samples. As expected, *Staphylococcus* incidence almost directly reflects the total number of species in the samples (rho above 0.8).

Regarding RT time, it displayed a significant moderate correlation with the tPSA value, negatively, which probably refers to the progression of therapy.

Univariate analysis of the parameters that significantly changed across time points had shown that blood parameters such as the count of the lymphocytes, monocytes, neutrophils, and erythrocytes, as well as the level of hemoglobin were decreasing during and shortly after radiotherapy and were returning to higher levels during the follow up (**Tab. 4**).

**Tab. 4.**
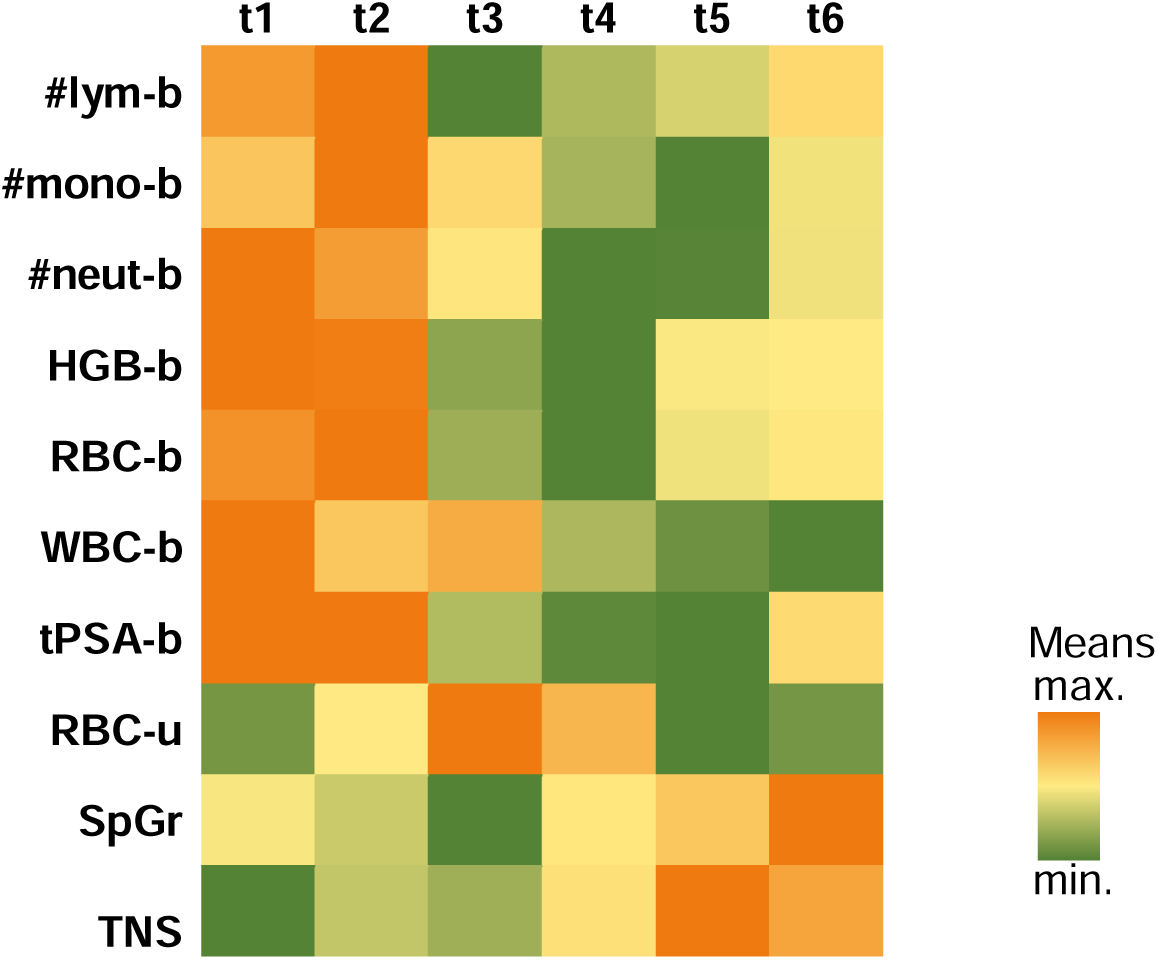
Summary heatmap, only showing variables that had a significant change in their means according to the analyzed time point (ANOVA-Tukey HSD, p adj. <0.05). t1 - before gold fiducial implantation into the prostate gland; t2 - before radiotherapy; t3 - at the end of radiotherapy; t4 - 1 month after radiotherapy; t5 - 4 months after radiotherapy; t6 - 7 months after radiotherapy. #lym-b – number of lymphocytes in blood, #mono-b – number of monocytes in blood, #neut-b – number of neutrocytes in blood, HGB-b – hemoglobin level in blood, RBC-b – erythrocytes level in blood, WBC-b – leukocytes level in blood, tPSA-b – prostate cancer specific antigen level in blood, RBC-u – erythrocytes level in urine, SpGr –specific gravity of urine, TNS – total number of species

On the contrary, the specific gravity of urine and TNS in urine increased after crossing the time point t3 (end of the RT). In the case of erythrocyte count in urine, there was a gradual increase until the end of RT and then a decrease to the initial value.

Finally, analysis of the blood and urine parameters that significantly changed under the presence of the most predominant microbial species/genus has been performed (**Fig. 5**). The obtained results indicated that *Corynebacterium* incidence implied in urine higher pH and lower specific gravity while the presence of *Enterococcus* related to lower cholesterol and higher IL-6 level in the blood as well as much higher white blood cells count in urine (over 300% increased). In turn, *Kocuria* presence was characteristic for later time points and implied a decrease in hemoglobin and cholesterol (total and non-HDL and LDL type) content in the blood and was accompanied by a lower erythrocytes count. *Micrococcus* incidence occurred in patient groups with higher triglycerides and hemoglobin levels in the blood. *Staphylococcus* presence was related to higher tPSA and PT and slightly lower PT index in blood.

**Fig. 5.**
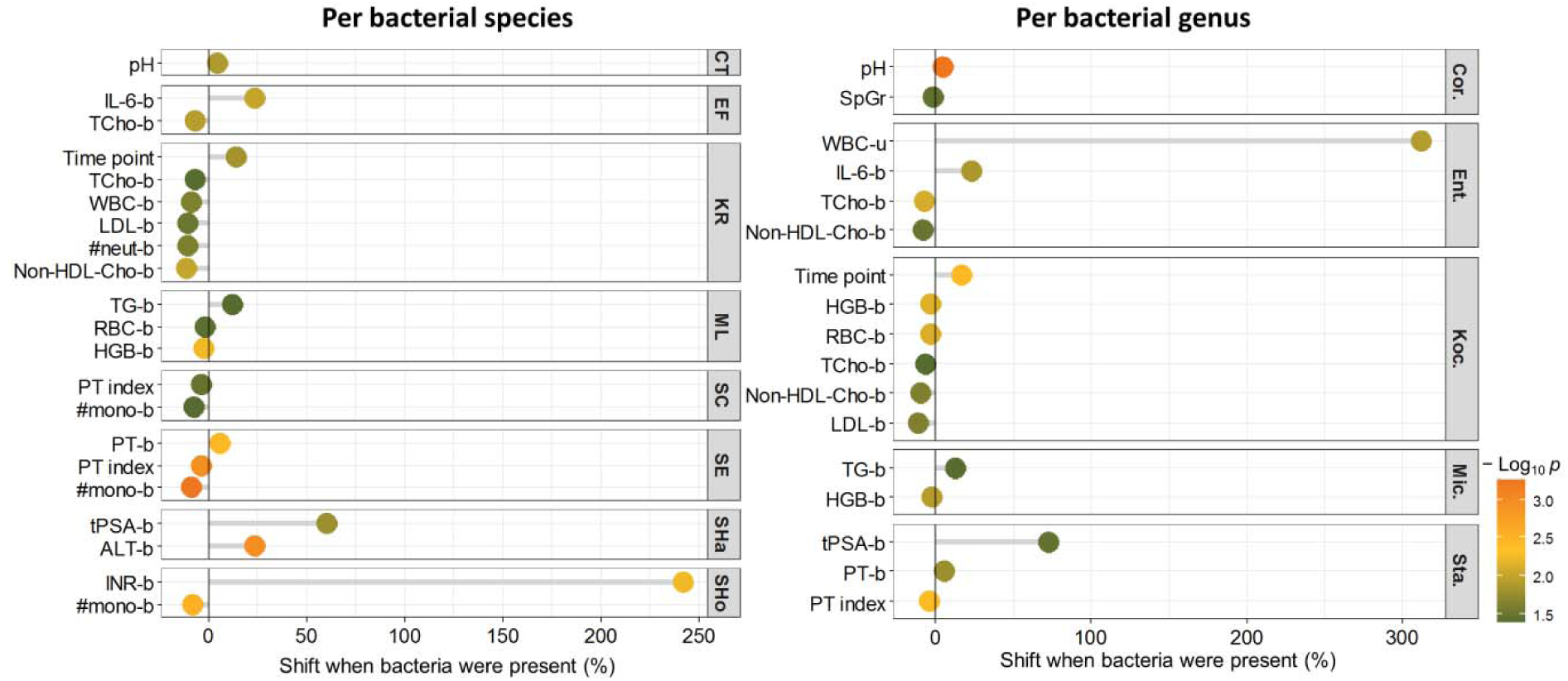
The results of the univariate analysis demonstrated significant changes in the value of the parameters depending on the presence of a given bacterial species/genus. Only bacteria displaying the most excellent frequencies among the samples were included. The presence or absence of bacteria in the samples was used as a categorical variable. Based on this classification, the means of numeric variables (i.e., biochemical parameters) were compared. Only significant comparisons (*t*-test, p<0.05) are shown. CT = *Corynebacterium tuberculostearicum*, EF = *Enterococcus faecalis*, KR = *Kocuria rhizophila*, ML = *Micrococcus luteus*, SC = *Staphylococcus capitis*, SE = *Staphylococcus epidermidis*, SHa = *Staphylococcus haemolyticus*, SHo = *Staphylococcus hominis*, Cor. = *Corynebacterium*, Ent. = *Enterococcus*, Koc. = *Kocuria*, Mic. = *Micrococcus*, Sta. = *Staphylococcus*

Concerning individual bacterial species, the observed higher pH of urine under the presence of *Corynebacterium* is related explicitly to the *C. tuberculostearicum* presence. The incidence of *E. faecalis* was most likely related to the higher IL-6 and lower total cholesterol in the blood. At the same time, *K. rhizophila* presence was responsible for all significant effects noted for *Kocuria* genus and additionally was related to the decrease in neutrophils count in blood. Regarding staphylococci, four species – *S. capitis*, *S. epidermidis*, *S. haemolyticus*, and *S. hominis*, have been recognized as having a significant relationship with the measured blood and urine parameters. Patients with *S. capitis* and *S. epidermidis* displayed lower PT index and monocyte counts. *S. haemolyticus* incidence was associated with patients displaying higher tPSA and ALT, whereas *S. hominis* incidence was accompanied by higher INR and lower monocyte count in the blood.

## 4. DISCUSSION

It was a long-held clinical dogma that urine was sterile until a series of recent studies demonstrated that urine contains microorganisms that are representative of a distinct flora in the urinary tract(^6^). Although there are recent hints in the literature pointing to the possible impact of the urine microbiome on prostate cancer malignancy, it is still virtually unexplored(^22^). It is believed that the urinary microbiome can influence cancer development by regulating pathogenic infections since many infectious agents could act as cofactors in carcinogenesis, while commensal bacteria may prevent outgrowth of pathogenic ones(^23^). Routine urine culture techniques are mainly used to isolate and identify pathogens involved in developing urinary tract infections (UTIs) since they enable the detection of fast-growing organisms like *E. coli* or *E. faecalis*. In contrast, the slow-growing ones are rarely isolated(^24,25^). Using 16S rRNA sequencing for microbial identification helps to overcome such limitations and therefore enables a more accurate investigation of the urinary tract microbiome. However, this approach suffers from another constraint, namely, it cannot differentiate between live and dead bacteria or bacterial DNA fragments. Hilt et al.(^25^) addressed this issue by expanding culture conditions, which resulted in a significant increase in the bacteria detection in samples previously classified as negative in terms of microbial presence – bacteria had grown in 80% of 65 samples of which 92% were culture negative using traditional culture techniques. Additionally, the authors revealed that most of the bacteria detected using RNA sequencing were cultivable under laboratory conditions. Similar to this study we used multiple culture conditions combined with the MALDI identification, which allowed us to decipher the rich urinary microbiota inhabiting the urinary tracts of the studied prostate cancer patients. We found over 1800 different microbial isolates representing 157 species and 45 genera. Identified microbiota comprised of both fast-growing common staphylococci, streptococci, enterococci or enterobacteria as well as those considered as rare or slow-growing, e.g. *Paracoccus* sp., *Roseomonas mucosa*, *Kytococcus sedentarius* or *Cutibacterium avidum*. Dubourg et al.(^26^) using culturomics approach (various culture conditions – 26 and MALDI TOF MS identification) isolated 450 different bacterial species, including 256 never described in urine and revealed that many members of the microbiota in the urinary tract are derived from the gut, thus, a paradigm shift is needed to better understand urinary microbiota composition. As we showed, such an approach is also valuable in tracking changes within urinary microbiota under different factors, e.g., radiation treatment.

The applied approach allowed us to indicate time points with the biggest microbiota changes (4 months after the end of RT), to determine the most critical biochemical parameters affecting the microbial composition of the urine samples (glucose level in the urine and blood) as well as to reveal the correlation of the presence of the certain microbial species with the patient’s biochemical parameters of the blood and urine. Regarding the last one, although significant shifts in the values under the presence of distinct bacterial species have been noted in the case of several parameters, most of them were low (a few percent). Nevertheless, the analysis showed that the presence of the *Enterococcus* spp., especially *E. faecalis*, was accompanied by significantly higher white blood cells count in urine (over 300% increased) and higher levels of IL-6 in the blood. It is known that many pathogenic microorganisms including opportunistic endogenous *Enterobacteriaceae* like *E. coli* or *Pseudomonas* spp., can infect the prostate and induce inflammatory response(^27^). In the case of *Enterococcus* spp., there is evidence of their pro-inflammatory role by i.a. inducing pro-inflammatory cytokines secretion such as IL-1β, IL-6 or TNF-α(^28–30^). Given this, the results of our analysis proved that the presence of *Enterococcus* demonstrates a negative effect on the healthy parameters of the patients; however, such an effect was not associated with the RT progression – nor *Enterococcus* count and IL-6 level correlated with time points.

Concerning shifts in the urinary microbiota diversity it is worthy to notice that among several measured biochemical parameters, the glucose level in the urine demonstrated the most remarkable correlation with the biodiversity of the urine microbiota – the higher the glucose content in urine, the higher the number of microbial species. Also, the glucose level in the blood contributed to this phenomenon, however, in a weaker manner. It is well documented that higher glucose concentrations in urine may promote the growth of bacteria(^31,32^). It was also shown that presence of the glucose in the urine stimulates biomass production of both gram-negative (e.g. *E. coli*, *P. aeruginosa*, *K. pneumoniae*) and gram-positive (e.g. *S. aureus* and *E. faecalis*) bacteria as well as increase their metabolic activity(^33^). Moreover, there are reports that high glucose concentrations impair epithelial barrier functions of the urinary tracts together with altering cell membrane proteins and cytoskeletal elements, resulting in increasing bacterial burden(^34,35^). Although most of the studies concerning the impact of the glucose level on the bacteria presence in the urine related to the UTIs development and diabetics, our study showed that such factors should be also considered during the analysis of the RT effect on the urinary microbiota. Increased glucose levels both in blood and urine are a factor contributing to the occurrence of infections. Moreover, increased glucose levels together with radiation-induced changes in the bladder wall, may increase the risk of developing infection and increasing radiation toxicity. In fact, in only 4 patients all glucose measurements were within the normal range, in the remaining patients at least one, and in many cases more than one, was above the norm. Additionally, several of our patients diagnosed with diabetes were taking medications that increased glucose excretion into urine, which, in light of our results, increases the risk of infection. Therefore, it is important to monitor its level during oncological treatment to minimize the risk of infection and properly diagnose and treat the diabetes and glucose intolerance.

While the primary goal of radiotherapy is to eliminate cancer cells, it is noteworthy that it may also have an impact on the balance of the microbiome. Research conducted thus far suggests a reciprocal relationship between radiotherapy and the microbiota. On one hand, the oxidative stress and inflammation caused by radiotherapy may disturb the microbiota, leading to a pro-inflammatory environment. Conversely, a disrupted microbiota has the potential to diminish the effectiveness of radiotherapy(^10^). Most of these observations were made concerning the gut microflora, which directly affects the functioning of the immune system(^36,37^). Matson et al., based on studies conducted on mice, observed that gut microflora can also migrate to other closely related tissues and influence cancer progression(^38^). Nevertheless, our understanding of the interplay between prostate cancer and the urogenital microbiome, particularly regarding how the microbiome influences the onset, progression, treatment response, and overall development of the disease, remains rather limited(^39^). Understanding these interactions is crucial for advancing personalized cancer treatments and optimizing patient outcomes. In our study, the presence of *Staphylococcus* was significantly associated with higher tPSA, and their presence was found in majority of samples, but it is still unknown, whether microbiome-based biomarkers can represent new diagnostic and prognostic factors.

Our study revealed that the urinary microbiome of prostate cancer patients was impacted by radiotherapy. Immediately after RT, less diversity of the microbiome was observed, which gradually increased over time after treatment (1-7 months). Moreover, its species composition was more diverse compared to samples before the implementation of treatment. The literature indicates that radiotherapy leads to dysregulation of microflora (especially intestinal), which is often manifested by a reduction in the number and diversity of intestinal microflora(^40^). It is generally accepted that radiation exposure contributes to sterilization(^41^). A significant increase in the diversity of microbial species within 1-7 months after finishing RT is most likely associated with a decline in the commensal bacteria counts noted right after the end of RT. It is believed they play a vital role in the homeostasis of the urinary tract, especially in terms of preventing pathogens’ development by outcompeting pathogens for shared resources, killing them by producing antimicrobial compounds, creating barrier-blocking pathogens access to the uroepithelium or priming immune defenses(^6^). It could be concluded that RT’s sterilization effect caused diminishing regulatory action of the urinary microbiota, facilitating the colonization of the urinary tract by various microorganisms. Pan et al.(^42^) hypothesized that RT may play a protective role against urinary tract infections in patients with prostate cancer since it may upregulate urine microbiome by reducing inflammations in the prostate via causing local immunosuppression, which may make it less likely for bacteria to cause urinary tract infections. Based on our results, this hypothesis could be valid for the time framework around the radiation therapy and directly after its finishing, where the RT sterilization effect was profound. Concerning a longer period after finishing RT, we can only conclude that the urinary tracts of prostate cancer patients are characterized by an increased propensity to colonization by microorganisms. Moreover, such conditions persist for a relatively long period - at least seven months after the end of RT. In contrast, the direction of the microbial shift (towards beneficial microbiota or pathogenic) is hard to predict since it is most likely dependent on the patient’s diet and risk of infection. Although the disturbed commensal microbiota and its protective role create a risk of infection, such conditions also allow a more straightforward introduction of beneficial probiotic bacteria. Therefore, patients undergoing RT should take special care of their supplementation, especially with probiotics. Such probiotic supplementation should not be postponed, since changes in the urinary microbiota of patients are visible as early as one month after the end of therapy, and the state of highest sensitivity to changes persists until the fourth month after the end of RT. Further studies should be conducted to evaluate the use of prebiotics and probiotics, medications, and dietary modifications in this group of patients to restore favourable bacterial flora. It is important to use the appropriate strain and understand its interaction with treatment. By comparing the gut microbiome of cancer patients who responded to immunotherapy with that of cancer patients who did not respond, it was found that the relative abundance of *Akkermansia muciniphila,* B*ifidobacterium longum, Collinsella aerofaciens* and *Enterococcus faecium* correlated with clinical response to this immunotherapy in cancer patients(^43,44^). *Akkermansia muciniphila* was shown to play a critical role in the reduction of abdominal IR-induced intestinal damage and application of probiotics or their regulators, such as metformin, could be an effective treatment for the protection of radiation exposure-damaged intestine(^45^). Metformin, a antidiabetic drug not only enhance beneficial bacteria, but also is known to enhance tumor response to radiation in experimental models, and retrospective analyses have shown that diabetic cancer patients treated with radiotherapy have better outcomes when they take metformin to control their diabetes(^46^).

It is also important to highlight that certain microorganisms exhibit resistance to elevated levels of ionizing radiation, which may contribute to the direction of the shift in the species composition within urinary microbiota. The survival and adaptation of bacteria to stressors involve intricate regulatory networks, encompassing post-transcriptional regulators like small RNAs. When effectively orchestrated, these mechanisms may bolster bacterial resilience to ionizing radiation(^47^). The least sensitive to ionizing radiation are the gram-positive strains *Deinococcus spp*. and *Rubrobacter spp*., which resist and survive doses of γ-ray greater than 25 kGy(^48^). Based on our research, the composition of the urine microbiome consists mainly of gram-positive strains, however, their percentage increases significantly after the end of RT. Deng et al. made similar observations, noting a higher incidence of gram-positive cocci and a lower incidence of gram-negative bacilli in patients who underwent radiotherapy for nasopharyngeal cancer compared to those treated with other therapies(^49^). In our research, gram-positive strains especially those belonging to the genus *Corynebacterium, Staphylococcus*, and *Streptococcus* were dominant during RT. Shrestha et al. noted a very similar composition of the urinary tract microflora in men and suggested that the urinary microbiome may influence chronic inflammation contributing to the development of prostate cancer(^50^). One of the first studies of the prostate microbiome conducted by Cavarretta and colleagues showed that *Staphylococcus* strains are more common in tumor and peri-tumor tissues(^51^).

*Kocuria rhizophila* emerged as a statistically significant strain, with its incidence showing a significant increase in patients post-radiotherapy(^52,53^). Similarly, although not statistically significant, similar observations were noted for strains of *Micrococcus luteus*. Probably, the higher capacity of these strains to endure harsh conditions, like radiation, might be attributed to the activation or expression of specific proteins in response to radiation-induced stress. The literature provides information about isolates of these species that exhibit increased resistance to radiation stress(^48,54^). Further research should focus on determining the microflora using molecular and spectrometric tests and linking it with the occurrence of urinary tract reactions, which may enable the selection of patients who are at higher risk of urinary tract complications after radiotherapy.

## 5. CONCLUSIONS

In conclusion, our study reveals that radiation therapy (RT) for prostate cancer induces a dynamic response in the urinary microbiome, characterized by an initial reduction in diversity post-RT followed by a subsequent increase. This augmentation in microbial diversity may be attributed to the sterilization effect of radiation, rendering the urinary tract more susceptible to colonization by various microorganisms in the RT-following period. The robustness of our methodology established in the previous work, employing MALDI identification alongside diverse culture conditions, has proven helpful in precisely tracking these intricate changes in the urinary microbiome during RT. Furthermore, our findings highlight the significant influence of glucose levels in both urine and blood on the urinary microbiota. To mitigate the potential pathogen burden after RT, considering the prolonged vulnerability of the urinary tract, proactive measures such as probiotic supplementation and careful management of glucose levels could be valuable strategies to maintain a healthier microbial balance. *Staphylococcus* presence was related to higher tPSA and its importance should be further investigated. These insights contribute to the evolving understanding of the interplay between RT, the urinary microbiome, and patient health, paving the way for more targeted interventions and personalized approaches in prostate cancer treatment.

## Acknowledgement

We would like to thank all participating patients. M.Z., E.S., P.F., and P.P., are members of Toruń Center of Excellence “Towards Personalized Medicine” operating under Excellence Initiative-Research University. A.A. is a member of Emerging Field “Cells as Experimental platforms and bioFACTories (CExFact)”.

## Authors contributions

Conceptualization, D.G., M.Z., P.P.; data curation, F.M., M.Z.; formal analysis, E.S., P.F., F.M., K.B.; funding acquisition, D.G.; investigation, E.S., P.F., W.M., K.K.; methodology, M.Z., P.P.; resources, D.G., P.P. .; supervision, M.Z.; validation, F.M., A.A.; writing— original draft preparation, E.S., M.Z., D.G.; writing—review and editing, D.G., P.P., R.K., M.R., E.T., J.M.-K. ; project administration, D.G., P.P. . All authors have read and agreed to the published version of the manuscript.

## Institutional Review Board Statement

The study was conducted in accordance with the Declaration of Helsinki and approved by the Institutional Ethics Committee of NIO-PIB Gliwice, Poland (protocol code KB/430-104/19 and date of approval 19 December 2019), and conducted in accordance with the principles of Good Clinical Practice.

## Data Availability Statement

Raw data used for the analysis and drawing conclusions are available on Repository for Open Data RepOD at https://doi.org/10.18150/KFIORE

